# Subcellular analyses of planarian meiosis implicates a novel, double-membraned vesiculation process in nuclear envelope breakdown

**DOI:** 10.1101/620609

**Authors:** Longhua Guo, Fengli Guo, Shasha Zhang, Kexi Yi, Melainia McClain, Claus-D. Kuhn, Tari Parmely, Alejandro Sánchez Alvarado

## Abstract

The cell nuclei of Ophisthokonts, the eukaryotic supergroup defined by fungi and metazoans, is remarkable in the constancy of both their double-membraned structure and protein composition. Such remarkable structural conservation underscores common and ancient evolutionary origins. Yet, the dynamics of disassembly and reassembly displayed by Ophisthokont nuclei vary extensively. Besides closed mitosis in fungi and open mitosis in some animals, little is known about the evolution of nuclear envelope break down (NEBD) during cell division. Here, we uncovered a novel form of NEBD in primary oocytes of the flatworm *Schmidtea mediterranea*. From zygotene to metaphase II, both nuclear envelope (NE) and peripheral endoplasmic reticulum (ER) expand notably in size, likely involving *de novo* membrane synthesis. 3-D electron microscopy reconstructions demonstrated that the NE transforms itself into numerous double-membraned vesicles similar in membrane architecture to NE doublets in mammalian oocytes after germinal vesicle breakdown. The vesicles are devoid of nuclear pore complexes and DNA, yet are loaded with nuclear proteins, including a planarian homologue of PIWI, a protein essential for the maintenance of stem cells in this and other organisms. Our data contribute a new model to the canonical view of NE dynamics and support that NEBD is an evolutionarily adaptable trait in multicellular organisms.

## Introduction

Nuclear envelope (NE), which is the boundary of the nucleus, is a defining feature of all eukaryotes. NE serves as a barrier for cytoplasmic and nuclear contents and activity, i.e., protein translation, mRNA transcription and DNA replication. Yet, it also poses a challenge to eukaryotic cell divisions: to separate linear chromosomes enclosed by the NE through assembly/disassembly of microtubules located in the cytoplasm.

Nature has evolved diverse solutions in Ophisthokonts to tackle this challenge of cell division [1–8]. Such solutions involve multiple modes of NE remodeling to allow accessibility to chromosomes by microtubules. The most straight-forward solution is open mitosis. As NE ruptures into pieces, chromosomes are completely exposed to cytoplasmic microtubules and establish contact through kinetochores. Cases were found, mostly in unicellular organisms, that complete rupture of NE is not necessary. In semi-open mitosis, small holes open locally on NE for adjacent microtubules to access condensed chromosomes within the nuclei. In closed mitosis, microtubule organization center (MTOC), is embedded in the NE during all or part of the cell cycle.

Among all the diverse modes of NE regulation during cell divisions, whether vesiculation is a disfavored strategy by natural selection remains controversial [9, 10]. The fate of NE proteins after NE breakdown and the source of NE proteins for the assembly of new NE in daughter cells underlies the motivation of a proposed vesiculation model four decades ago [11, 12]. In this model, the nucleus breaks down into multiple vesicles with pieces of NE enclosing portions of the nuclear content, while chromosomes are exposed to cytoplasmic factors. Accumulating evidence supports an otherwise mutually exclusive model, that NE proteins are dispersed into the peripheral ER upon NEBD and comes from the ER network upon assembly of a new nucleus, and challenges the experimental methods in earlier studies. While in principle, NE vesiculation maintains barrier function between cytoplasm and nucleus materials, as is in closed mitosis, and allows for full accessibility to the condensed chromosomes by microtubules, as is in open mitosis, whether this solution for cell division indeed exists in nature needs direct evidence.

## Results and Discussion

### Meiotic progression can be detected and stages quantified in planarian ovaries

Here, we examined NEBD during oocyte meiosis in a free-living fresh water flatworm, *Schmidtea mediterranea*, which has been established as a model system to study adult stem cells, regeneration, and germ cell specification [13–22]. Detected widespread maintenance of genome heterozygosity suggests potential mechanisms in meiosis [23, 24]. Yet, meiosis has only been studied in the testis [25].

To characterize female meiosis in *S. mediterranea*, we examined the ovaries using Transmission Electron Microscopy (TEM). Ultrastructural studies revealed five categories of cells with oocyte features (Figure 1; Supplementary Fig.1). These cells are in close proximity to each other in a relatively compacted area of the ovary (Figure 1A), are of much larger size (20~50μm in diameter) than most somatic cells (10~20μm), and contain germ cell specific organelles (*e.g*., chromatoid body [26–35] and annulate lamella [36–43]) (Supplementary Fig.1). We grouped cells into five categories based on their nuclear morphology. Type-I cells have smaller nuclei with multiple Synaptonemal Complexes (SYCPs) [44–47] (Figure 1B, Supplementary Fig.2), suggesting they are oocytes at zygotene or pachytene stage of prophase I. Type-II cells have undulating NEs, and remnants of SYCPs, characterized by high electron density, short dark stripes (Figure 1B). The dissolution of SYCP suggests Type-II cells are entering diplotene stage of prophase I. Type-III, IV and V cells have numerous vesicular structures surrounding the NE and dense patches of condensed chromatins inside the NE (Figure 1C-E). In Type-IV cells, vesicles are elongated. In Type-V cells, the vesicles are in the cytoplasmic periphery, and shaped like dumbbells.

**Figure 1.**
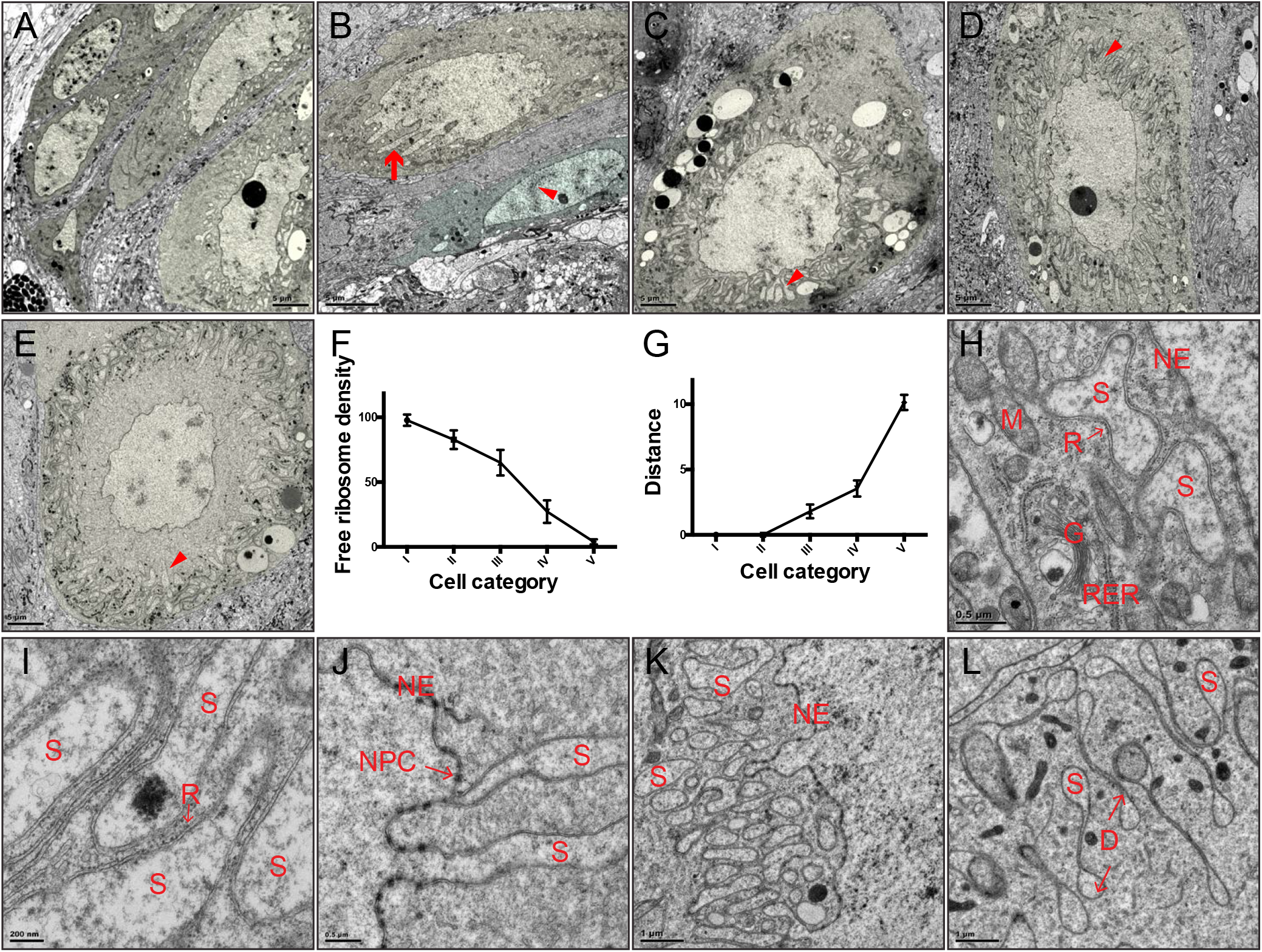
Ultrastructural studies of oocytes at prophase I in *S. mediterranea*. (A) A section of the planarian ovary. Oocytes were pseudo-colored in light yellow. (B) Type I (pseudo-colored in light green) and Type II (pseudo-colored in light yellow) cells. Arrow=undulation; arrowhead=SYCP. (C) Type III cells (pseudo-colored in light yellow). Arrowhead=vesicles. (D) Type IV cells (pseudo-colored in light yellow). Arrowhead=vesicles. (E) Type V cells (pseudo-colored in light yellow). Arrowhead=vesicles. (F) Free ribosome densities in the cytoplasm (number of free ribosomes per 400nm by 400nm area). Three oocytes were quantified for Type I to Type IV cells. Four oocytes were quantified for Type V cells. For every oocyte, three distant areas were quantified. (G) Distances between distal end of the vesicles to the nuclear envelope. Six oocytes for Type III, four oocytes for Type IV, and three oocytes for Type V cells were quantified. Distance unit is μm. (H) Perinuclear vesicles have double membranes. NE=nuclear envelope. M=mitochondria. R=ribosome. G=Golgi. RER=rough ER. S=perinuclear vesicles. (I) Ribosomes decorate the outer membrane of perinuclear vesicles. S=perinuclear vesicles. R=ribosomes. (J) Perinuclear vesicles can directly connect with nuclear envelope. NPC=nuclear pore complex. S=perinuclear vesicles. NE=nuclear envelope. (K) Perinuclear vesicles are dumpy and close to nuclear envelope in Type III cells. (L) Perinuclear vesicles are in dumbbell shapes in Type V cells. D=dumbbell.

As free ribosomes are easily recognizable and almost evenly distributed in the cytoplasm of all cells, we quantified densities of free ribosomes to examine relationships of these cells. From Type-I to Type-V cells, a gradual decrease in free ribosome density was observed, suggesting Type-I to Type-V cells are oocytes at progressive steps of meiosis (Figure 1F). Consistently, distances of the proximal ends of the vesicles to the NE steadily increase from Type-III to Type-V cells (Figure 1G), which are diplotene to diakinesis stages of prophase I. As ovulated oocytes are arrested at metaphase II [23], meiosis stages from prophase I to metaphase II likely take place as the oocytes travel through the tuba and oviduct to the female atrium [48]. Alternatively, missing steps in meiosis could be fast and transient, which would be difficult to detect in the ovary.

### Nuclear envelope breakdown of planarian oocytes yields abundant, double-membraned vesicles

Five features define the perinuclear vesicles as novel NE-associated subcellular organelles. First, they are double membraned and distinct from peripheral ER (Figure1H). Second, ribosomes decorate the outer membrane of the vesicles (Figure1H,I). Third, electron density in the interior of the vesicles are comparable to nucleoplasm, but distinct from cytoplasm (Figure1H,J,K). Fourth, distances between the inner and outer membranes are comparable to the NE (Figure1H,J,K). Fifth, membranes of some vesicles can be physically continuous with the NE but lacking nuclear pore complexes (NPCs) (Figure 1J). Interestingly, NPCs are present in the NE immediately adjacent to the emerging vesicles. Specific regulations of NPCs on the vesicles are likely due to three mechanisms: vesicles form with newly synthesized NE without NPCs; vesicles form with pre-existing NE on which NPCs are selectively disassembled; NPCs are disassembled and dispersed on the vesicle membranes.

While in Type-III cells the vesicles are dumpy and adjacent to the NE (Figure 1K), they are dumbbell shaped and far away from the NE in Type-V cells (Figure 1L). Hence, the formation of perinuclear vesicles is very dynamic. As it appears that these vesicles radiate from NE (Figure 1C-E), we named them Sunburst NE Vesicles (SNEVs).

### Nuclear membrane vesiculation products are topologically complex

To clarify the dynamics of SNEV formation, we reconstructed 3-D models from serial sections of the oocytes. SNEVs start as double-membraned buds of NE in Type-III cells (Figure2A). The buds grow distally, branch out and fold onto themselves (Figure2B,). In Type-V cells, the SNEVs are elongated, with the proximal ends arranged as tubules and the distal ends as flattened, stacked sheets (Figure2C-E). Some SNEVs appear disconnected from the NE (Figure2C,F-I).

**Figure 2.**
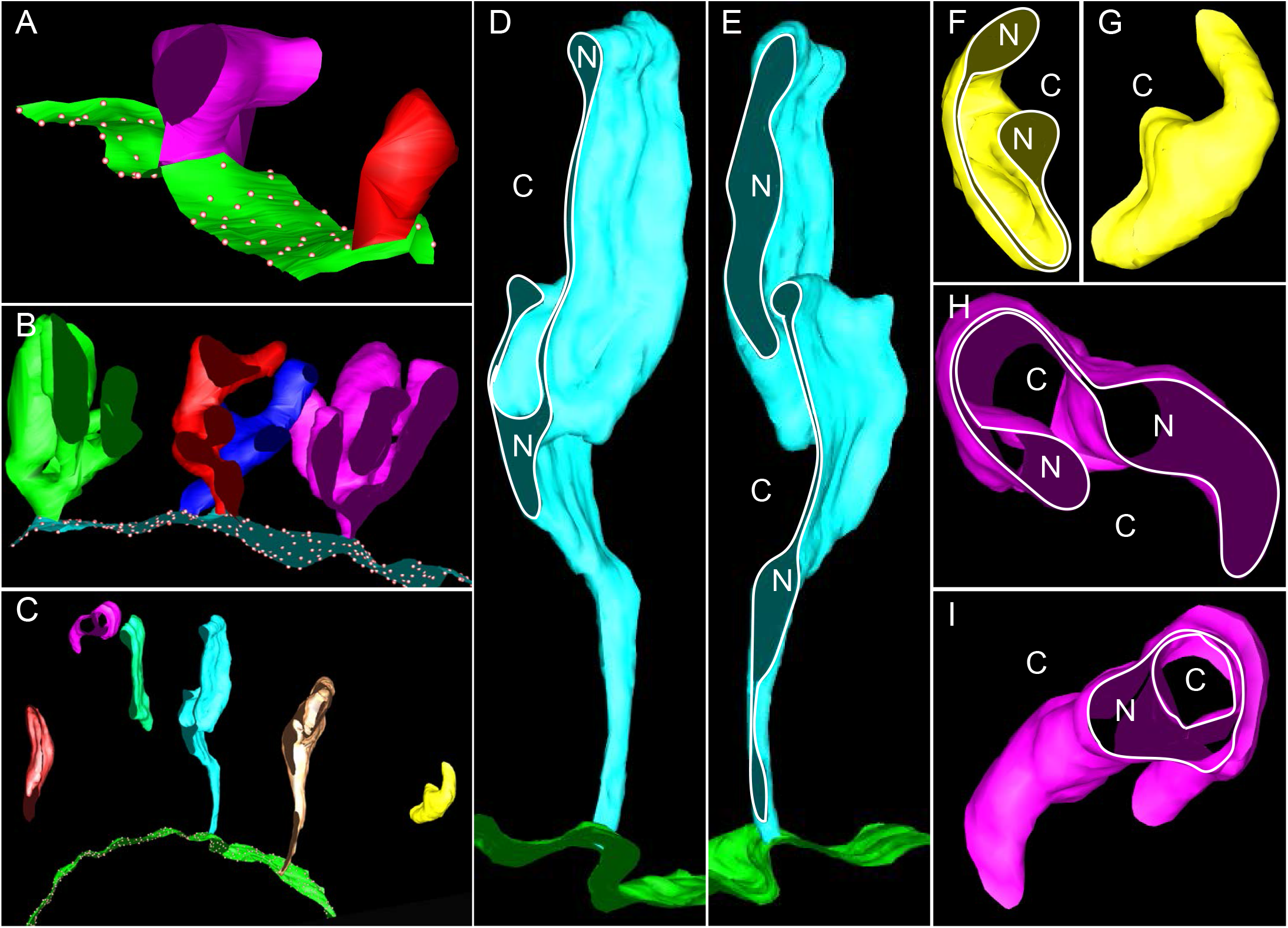
3D reconstruction of the Sunburst Nuclear Envelope Vesicles. (A) Type III cells. Green sheet=NE. Dots=NPC. Purple and red buds=SNEVs. (B) Type IV cells. Cyan sheet=NE. Dots=NPC. Green, Red, Blue, Purple coral shapes=SNEVs. (C) Type V cells. Green sheet=NE. Dots=NPC. The rest=SNEVs. (D-E) The same SNEV sectioned at different positions. N=nucleoplasm. C=cytoplasm. Ribosomes on NE (green sheet) were present but not illustrated. (F-G) The same SNEV viewed from different angles. N=nucleoplasm. C=cytoplasm. Dumbbell shapes in F. (H-I) The same SNEV sectioned at different positions. N=nucleoplasm. C=cytoplasm. Dumbbell shapes in H. Double rings in I.

3-D models revealed topological complexity of the SNEVs in Type-V cells. SNEVs are dumbbell-shaped in some sections, but appear with variable shapes (*e.g*., rings) in other sections (Figure2D-I). The encapsulated space of the vesicles is not spherical. Instead, some areas of the inner vesicle membranes are in close proximity.

To examine the fate of the SNEVs after oocyte maturation, ovulated oocytes in egg capsules, which are arrested at metaphase II [23], were studied. In general, cytoplasmic space of ovulated oocytes is filled in its entirety with membrane units of variable sizes and shapes (Figure3A-B). Nonetheless, all membrane units are topologically similarly organized (Figure 3B). Switching from short and oval shapes of the SNEVs in Type-III oocytes to dumbbell shapes in Type-V oocytes implicates a tendency of double-layered membranes to form four-layered doublets (Figure 3C, left to right). The membrane units in metaphase II oocytes have long stretches of such four-layered doublets (Figure3D). In fact, these four-layered doublets are very prominent and comparable to NE doublets in mice and human oocytes after germinal vesicle breakdown (GVBD) [49–51] (Supplementary Figure 3D). In both cases, the inner membranes of the two double layered membranes are the contacting surfaces. Doublet formation could be a general property of membranes in the oocytes.

**Figure 3.**
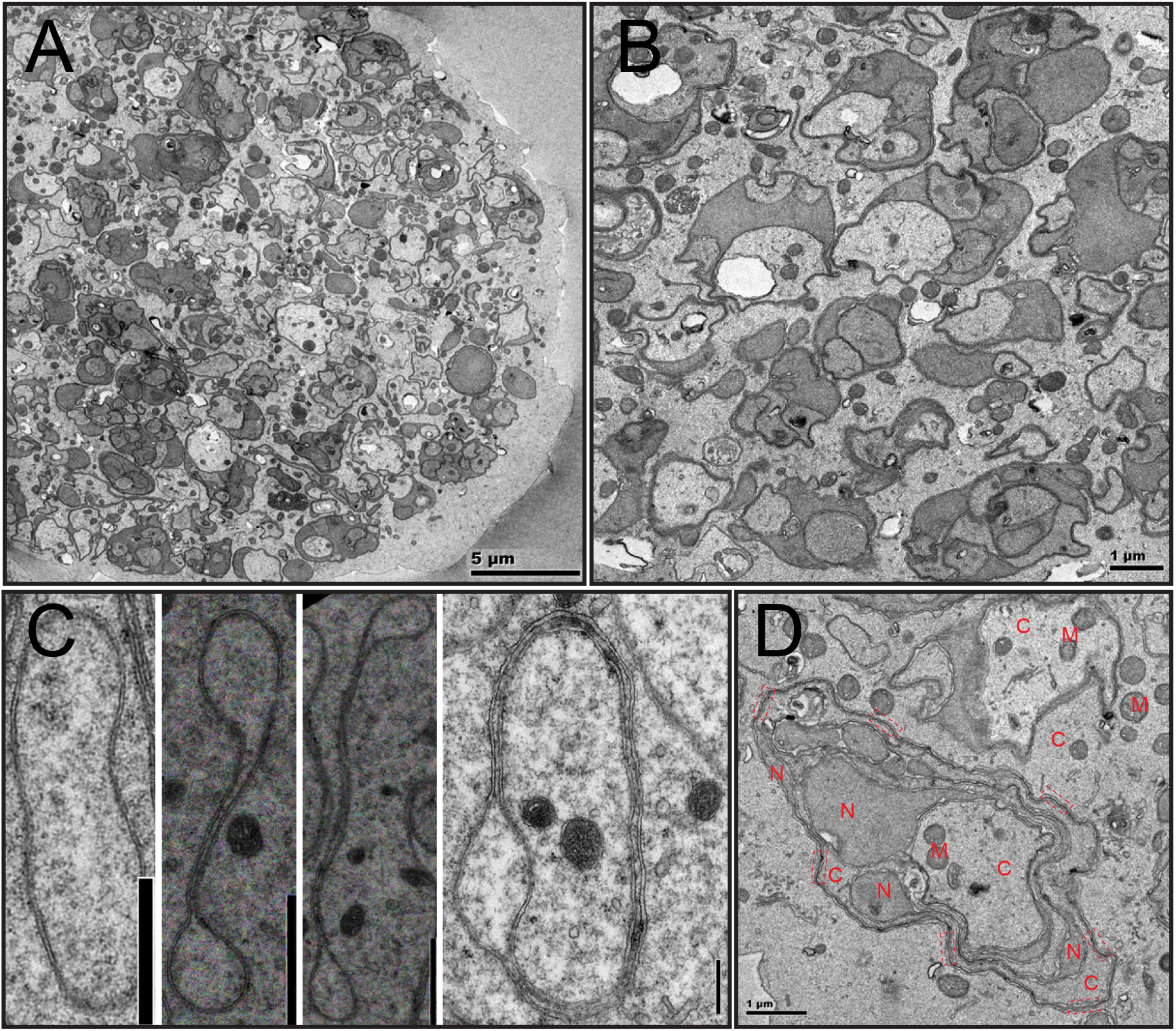
Ultrastructural view of ovulated unfertilized oocytes at metaphase II. (A) Overview of an unfertilized oocyte at metaphase II. (B) SNEV units in the cytoplasm with various sizes and shapes. (C) From dumpy SNEVs in Type III cells (left), to SNEVs in Type V cells (right), the inner membranes tend to adhere to form NE doublets. Scale bar=1 μm. (D) SNEV units in metaphase II oocytes have complex organizations, and prominent NE doublets. Dashed line rectangles=NE doublets. C=cytoplasm. N=nucleoplasm. M=mitochondria.

Taken together, the dynamics of SNEVs and the disappearance of nuclei in metaphase II oocytes define a novel form of NEBD or GVBD.

### Double-membraned vesicles are filled with nucleoplasmic proteins

To examine the fate of nuclear proteins during NEBD, antibodies against the Argonaute protein family PIWI protein SMEDWI-2 [52–56] and Histone H3 were used to characterize the dynamics of SNEV formation. Immunohistological studies revealed SMEDWI-2 persists in the nucleus during all stages of prophase I, where it marks SNEV-like structures (Figure4A-C). This is contrast to cytoplasmic SMEDWI-1 protein, which is degraded as the oocyte matures (Supplementary Figure 3A). Histone H3 protein shows the same dynamics in the nucleus and in SNEVs as SMEDWI-2 (Supplementary Figures 3B-C). These data conclude nuclear proteins are packaged into SNEVs. Interestingly, chromosomes are specifically excluded since these vesicles are negative for DNA dyes (*e.g*., DAPI, Hoechst 33342) (Figure 4, Supplementary Figures 3B-C).

**Figure 4.**
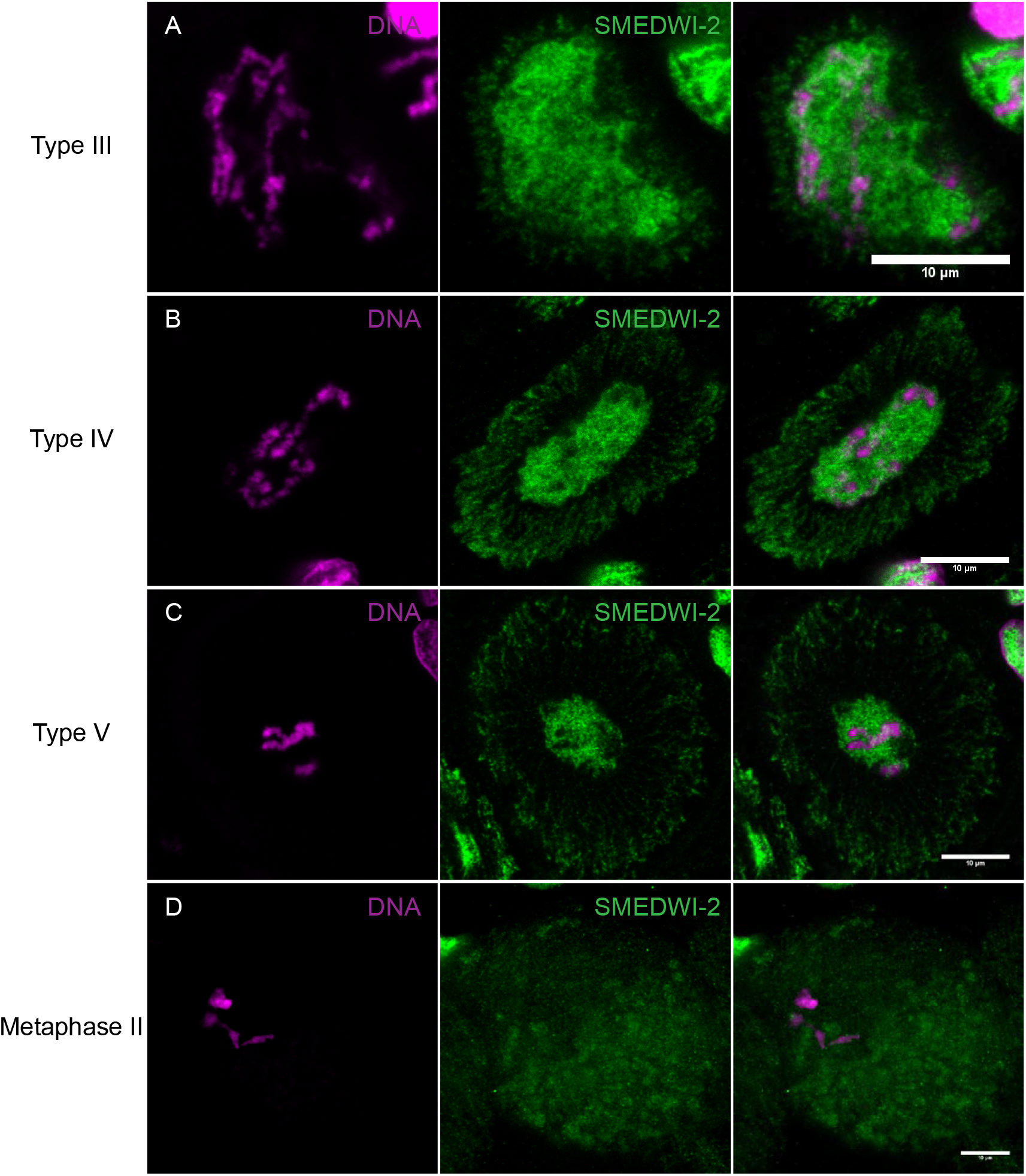
Nuclear contents (*e.g*. proteins) packaged and transported by SNEVs into cytoplasm. Planarian oocytes stained with SMEDWI-2 antibody. Left column=chromosomes with Hoechst 33342 staining. Middle column=SMEDWI-2 antibody. Right column=merge. (A) Type III cells. (B) Type IV cells. (C) Type V cells. (D) Rare metaphase II cells in the ovary.

Metaphase II stage oocytes with four condensed chromosomes can be found in the ovary at low frequency. The distribution of SMEDWI-2 protein in metaphase II stage oocytes (Figure 4D) in the ovary is consistent with our ultrastructural findings in the ovulated oocytes. There, the nucleus is dissolved into individual SNEV units of variable size and morphology (Figure 3), supporting the view that SNEV formation involves mass encapsulation of nuclear contents as vesicles dispersed throughout the cytoplasm.

### Marked expansion of nuclear double-layered membranes occurs during planarian oocyte NE breakdown

The formation of SNEVs is accompanied by an expansion of NE surface area. As oocytes mature, cell volume increases approximately 8 times from pachytene to diplotene stages (Figure4A,C; Figure5A-B; Supplementary Figure3B-C). To maintain a constant nuclear to cytoplasmic ratio [57–59], nuclear volume thus increases, leading to a 4-fold expansion of the NE surface area in case of a spherical nucleus. In addition, partitioning nuclear volume into much smaller vesicles leads to a significant increase in surface area. The larger the number of SNEVs, the more the membrane surface area increases. We estimated a 40-fold expansion of nuclear double-layered membranes in total (Methods).

**Figure 5.**
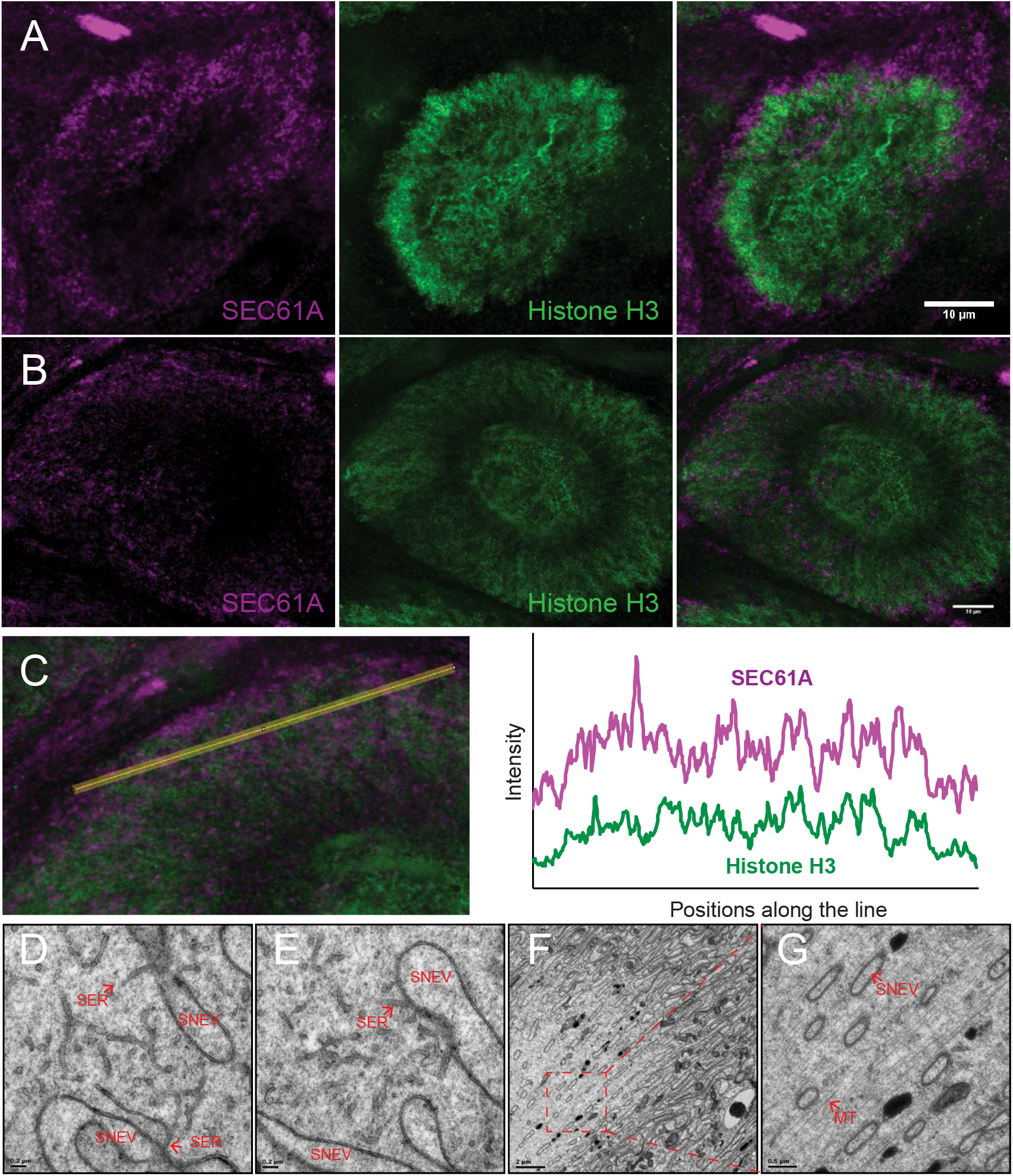
Interactions between SNEVs and peripheral ER, and microtubules. (A-C) Double staining with anti-SEC61A (magenta) and anti-Histone H3 (green) in Type III oocytes (A), and Type V oocytes (B and C). (C) Line profiling to examine co-localizations between peripheral ER (SEC61A, magenta), and SNEVs (Histone H3, green). (D-E) TEM view of peripheral smooth ER (SER) and SNEVs. (F-G) TEM view of microtubules (MT) and SNEVs.

### ER is unlikely to serve as a membrane reservoir for double-membraned vesicle biogenesis

To examine whether expansion of NE/SNEV surface area leads to a net decrease of peripheral ER, a SEC61A [60–64] antibody was used to visualize the ER network. The distinctive staining patterns of SEC61A and Histone H3 (Figure5A-B) suggest that SEC61A is mostly localized on the peripheral ER instead of the NE. Line profiling showed that SEC61A can co-localize with Histone H3 (Figure5B-C). However, even in such areas, most of the SEC61A signal is separated from Histone H3 signal in “salt and pepper” patterns (Figures5B-C). The independent organization and occasional physical interaction of peripheral ER and elongating SNEVs were verified by TEM (Figures5D-E). Direct interaction between tubular ER and elongating SNEVs suggests tubular ER may contribute to the elongation process. One possible function is to provide membranes. However, no clear reduction of the peripheral ER network is observed (Figures5A-B). Collectively, the ER-NE-SNEV membrane system expands in size as the oocytes mature. Hence, *de novo* membrane synthesis is likely required for the formation of SNEVs.

### Conclusions

Our data provided direct evidence that nuclear envelope can break down into vesicles during cell division, highlighting a new paradigm of nuclei dynamics in Ophisthokont. NE vesiculation is likely a trait adapted to the biology of the superphylum Platyhelminths. Establishment of double-membraned vesicles from NE were noted in female gonads in three other species from Platyhelminths, *Cura foremanii, Sabussowia dioica*, and *Vorticeros luteum* [65–67]. Confusion of these double-membraned vesicles with peripheral ER [68] only emphasizes that NE vesiculation has not been recognized.

What are the functions of SNEVs? Requirement of *de novo* membrane synthesis and specific regulation of NPCs support that SNEVs are not units for waste disposal but instead tightly regulated structures. As NE vesiculation was found specific to female meiosis, we speculate that SNEV formation is a strategy to regulate the fate of nucleoplasm after GVBD and the establishment of pluripotency in the zygote. While SMEDWI-1 is degraded during oocyte maturation (Supplementary Figure 3A), nuclear SMEDWI-2 is preserved. Importantly, loss of SMEDWI-1 does not show a phenotype in adult planarians, whereas abrogation of SMEDWI-2 leads to loss of somatic stem cells and death of the worms [53, 69–71]. Additionally, SNEVs may direct nucleoplasm to chromosomes for reassembly of the zygote nucleus, and jump-start mitotic divisions in early embryonic development.

## Materials and Methods

### Worm care

Planarians were maintained in 1x Milli-Q standard planarian medium at 18°C, with constant once or twice a week feeding of organic liver paste [72, 73]. To study the ovaries, sexually mature worms of 1 to 2cm in length were used. Multiple planarian lines were used for the analysis. Data reported were from line S2-3 and S2F8b [23] of *S. mediterranea*. To obtain ovulated oocytes, worms were maintained in solitude as virgins with twice a week feeding.

### Transmission Electron Microscopy

For TEM analysis, dissected ovaries were fixed with 2.5% paraformaldehyde/2% glutaraldehyde/PBS for overnight at 4°C. Then the tissues were processed as described with modifications [74–76]. Briefly, the tissues were washed with 0.1 M sodium cacodylate buffer (pH=6.8), intensified in 2% OsO4/0.1 M sodium cacodylate buffer (pH=6.8), stained with 2% uranium acetate *en bloc*, dehydrated with a graded ethanol series (30%, 50%, 70%, 95%, and two times 100%, 10 min each), equilibrated with two incubations (10 min) in propylene oxide, and incubated in 50% propylene oxide/50% Epon resin (EMS, Fort Washington, PA) mixture overnight. The samples were then infiltrated in 100% Epon resin for 4hr, embedded, and polymerized at 60°C for 24 hrs. After sectioning, the images were acquired on a FEI transmission electron microscope (Tecnai Bio-TWIN 12, FEI). 3D EM models were constructed using the IMOD image-processing package [77]. EM images were converted into stacks as .mrc files and then aligned using MIDAS. Volume segmentation, 3d meshing and surface rendering were done in 3dmod. ImageJ was then used for image format conversion.

Taking these two factors into account, we measured total circumference of SNEV membranes and NE membranes in Type-V and Type-I oocytes. Images used for the measurements were from sections of oocytes imaged by TEM. From this crude quantification,

### Histological sections

Sexually mature worms were fixed with freshly prepared 4% paraformaldehyde (Electron Microscopy Sciences, Catalog no.: 15710) in PBS for one hour at room temperature with gentle shaking. The anterior fragment of the planarians with the ovaries were obtained for paraffin or cryo sections after washing with 1x PBS for three times. The rinse was 20min each. For paraffin processing, worms are dehydrated through graded ethanol (30%, 50% in PBS) and then stored in 70% ethanol at 4°C for overnight. Paraffin blocks were loaded into a Tissue-Tek VIP processor (Sakura, Netherlands), followed by graded ethanol dehydration (70% for 15 min, 80% for 20min, 95% for 15min and 100% for 20min) and xylene substitute substance clearance (10 min/changes for 3 times). After 4 changes of paraffin infiltration (20 min/change), planarian was embedded for sectioning. Paraffin sections with 8um thickness were cut using a Leica RM2255 microtome (Leica Biosystems Inc. Buffalo Grove, IL) and mounted on Superfrost Plus microscope slides (Fisher Scientific,). For cryo processing, fixed planarian was dehydrated through 30% sucrose and followed by embedding with OCT compound (Tissue-Tek, CA). Cryo sections with 14um thickness were cut using a Leica CM3050S cryostat (Leica Biosystems Inc. Buffalo Grove, IL).

### Immunofluorescence staining

Paraffin or cryo sections of planarian fragments containing ovaries were used for immunofluorescence staining. Antibodies anti-SEC61A, anti-Histone H3 and anti-ds DNA was from Abcam (Catalog no.: ab183046, ab1791, ab24834, and ab27156). SEC61A is an evolutionarily conserved subunit of the Sec61/SecY complex, an ER apparatus that translocates nascent membrane proteins into the ER. SMED-SEC61A protein sequence is 86% identical to the human SEC61A isoform 1.Anti-SMEDWI1 was a kind gift from Dr. Jochen Rink. Anti-SMEDWI2 was a kind gift from Dr. Claus-D. Kuhn and Dr. Qing Jing. Goat anti-mouse IgG secondary antibody Alexa Fluor 488, and goat anti-rabbit secondary antibody Alexa Fluor 647 were from Thermo Fisher Scientific (Catalog no.: A-11001, and A-21245). Generally, histological sections were rinsed with PBS with 0.5% Triton X-100 for three times. The rinse is 10 minutes each. Tissues were digested with 2μg/ml Proteinase K (Thermo Fisher Scientific, 25530049) and 0.1% SDS for 10min at room temperature in PBS with 0.5% Triton X-100. After extensive washes, tissues were incubated with 10% Horse serum (Sigma, H1138) in PBS with 0.5% Triton X-100 for one hour at room temperature. All primary antibodies were used as 1:100 dilution in the blocking solution (10% Horse serum in PBS with 0.5% Triton X-100). Tissues were incubated with primary antibodies overnight at 4°C with gentle shaking. After three washes, the tissues were incubated with secondary antibodies overnight 4°C with gentle shaking. All secondary antibodies were used as 1:300 dilution in the blocking solution. Hoechst 33342 (Thermo Fisher Scientific, H3570) was used as 1:300 to stain the tissues for 30min at room temperature during washes. Slides were mounted with Prolong Diamond Antifade Mountant (Thermo Fisher Scientific, P36965).

### Image acquisition

All fluorescence images were acquired with ZEN software. All raw data were saved as 16bit images. Zeiss LSM-780 Confocal Microscope and Alpha Plan-Apochromat 100x/1.46 Oil DIC objective were used for most images reported. Zeiss LSM-710 Confocal Microscope and Alpha Plan-Apochromat 63x/1.46 Oil Korr M27 objective were used for Supplementary Figure 3A with a zooming factor of 0.7. For double staining with anti-SEC61A and anti-Histone H3 (Figure 5), lasers 633 and 488 were used. Fiji is used for all image processing.

## Acknowledgements

The authors would like to thank Drs. Jochen Rink and Qing Jing for sharing SMEDWI-1 and SMEDWI-2 antibodies. We would like to thank Dr. Elizabeth M. Duncan for suggestions and sharing of histone antibodies, Dr. Sue Jaspersen for revising the manuscript, and suggestions of ER markers, Drs. Christopher P. Arnold and Stephanie Nowotarski for revising the manuscript, Dr. Yongfu Wang for histological sections and antibody testing, Dr. Zulin Yu for assistance in imaging, Mr. Eric Ross for identifying reticulon homologues in *S. mediterranea*, and Mr. Mark Miller for illustration. We would like to thank Dr. Isabel Espinosa Medina for translation of the French article, Jillian Thaden, Katie Evans, Andrew Vogelsang, Corey Abrams, Shane M. Merryman, and Diana Baumann from the Stowers Aquatics Facility for skillful management of the planarians. This work was funded in part by NIH R37GM057260 to A.S.A. A.S.A is an investigator of the Howard Hughes Medical Institute and the Stowers Institute for Medical Research.

## Author Contributions

The project was conceived and designed by L.G. and A.S.A. Ultrastructural sample preparation and data collection were by F.G., L.G., K.Y. and M.M. Gene cloning, RNAi and phenotyping were by S.Z. and L.G. Figures were developed by L.G. Data interpretation and manuscript preparation were by L.G. and A.S.A.

---

**Supplementary Figure 1.**
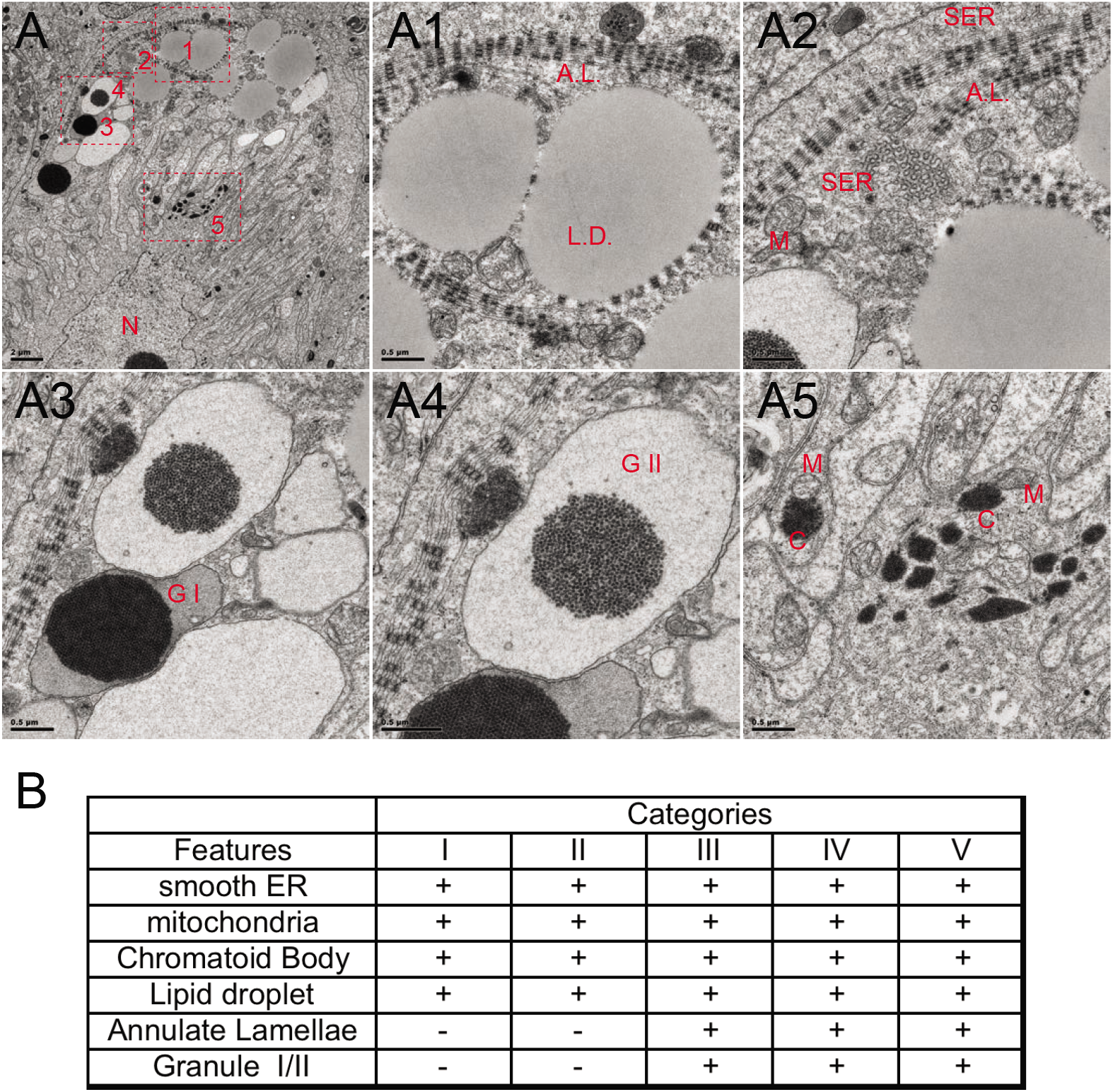
Characteristic features of a female germ cell. (A) Overview of a corner of a Type IV oocyte, and five regions which will be zoomed in from a TEM image. N=nuclear. (A1) A.L.=annulate lamellae. L.D.=lipid droplet. (A2) SER=smooth ER. A.L.=annulate lamellae. M=mitochondria. (A3) GI=Cortical granules, type I. (A4) GII=Cortical granules, type II. (A5) M=mitochondria. C=chromatoid body. (B) Summary of the presence and absence of all organelles in different types of cells.

**Supplementary Figure 2.**
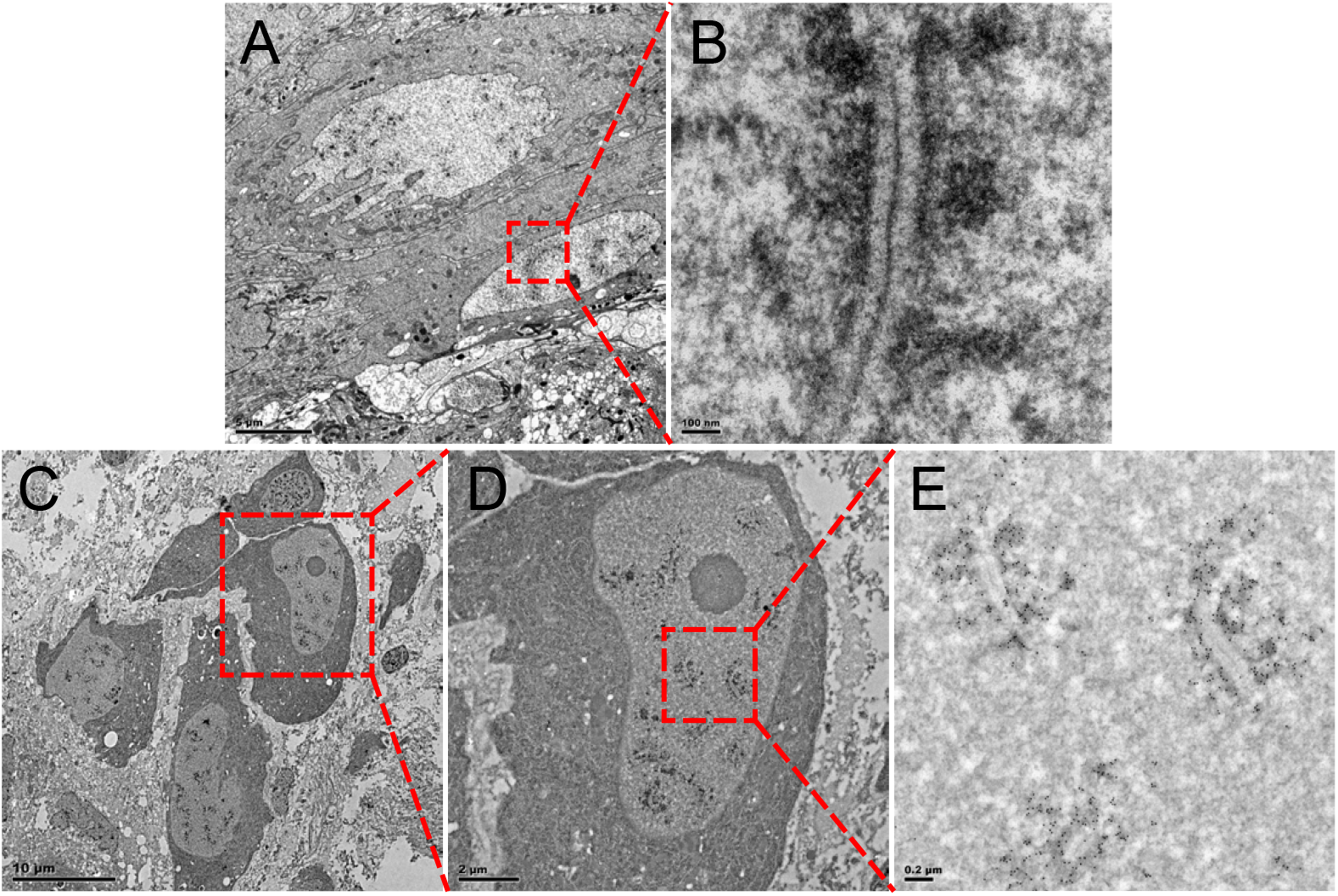
Type I cells have Synaptonemal Complexes. (A) TEM images of Type I (middle right) and Type II (top) cells. (B) Synaptonemal Complexes. (C-E) Immuno TEM studies of the Type I cells with anti-ds DNA antibodies. High electron density regions in the Synaptonemal Complexes are dsDNA (E).

**Supplementary Figure 3.**
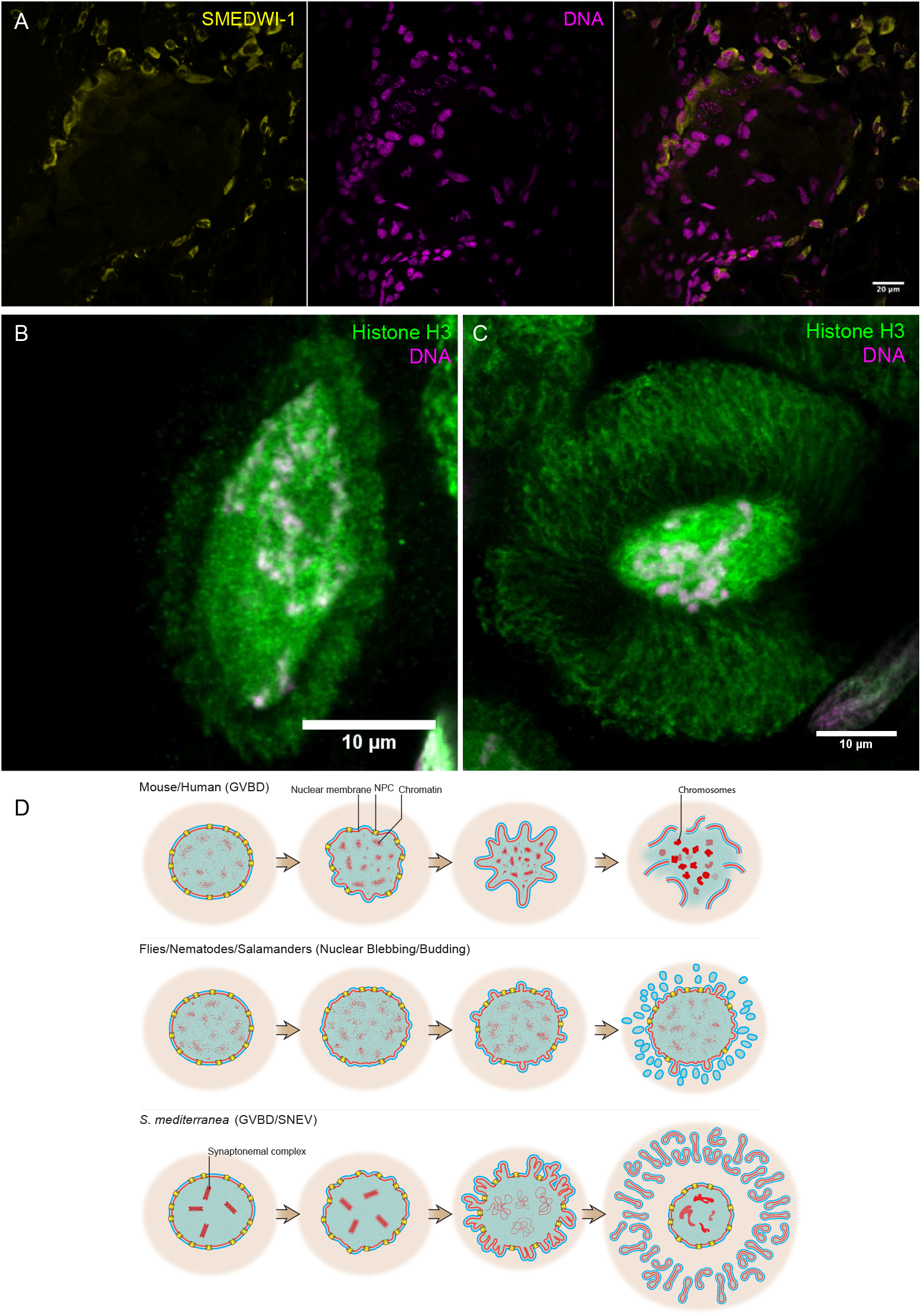
Immunofluorescence staining with SMEDWI-1 and Histone H3 antibodies. (A) A section of a planarian ovary stained with SMEDWI-1 antibody. Left=anti-SMEDWI-1. Middle=Hoechst 33342. Right=merge. (B-C) Planarian oocytes stained with Anti-Histone H3 antibody. Anti-Histone H3 (green) in Type III cells (B), and Type V cells (C). DNA is in magenta. Overlap of DNA and Histone H3 is white in color. (D) Schematic illustration of GVBD in mouse and human oocytes (top), nuclear budding/blebbing in somatic cells or oocytes of flies, nematodes, and salamanders (middle), and GVBD to produce SNEVs in the oocytes of *S. mediterranea*.

